# Organizational principles of the primate cerebral cortex at the single-cell level

**DOI:** 10.1101/2024.07.05.602270

**Authors:** Renrui Chen, Pengxing Nie, Liangxiao Ma, Guang-Zhong Wang

## Abstract

The primate cerebral cortex, the major organ for cognition, consists of an immense number of neurons. However, the organizational principles governing these neurons remain unclear. By accessing the single-cell spatial transcriptome of over 25 million neuron cells across the entire macaque cortex, we discovered that the distribution of neurons within cortical layers is highly non-random. Strikingly, over three-quarters of these neurons are located in distinct neuronal clusters. Within these clusters, different cell types tend to collaborate rather than function independently. Typically, excitatory neuron clusters mainly consist of excitatory-excitatory combinations, while inhibitory clusters primarily contain excitatory-inhibitory combinations. Both cluster types have roughly equal numbers of neurons in each layer. Importantly, most excitatory and inhibitory neuron clusters form spatial partnerships, indicating a balanced local neuronal network and correlating with specific functional regions. These findings suggest that different brain regions of the primate cortex may exhibit similar mechanisms at the neuronal population level.

## Main

Single-cell sequencing provides a profound opportunity to explore the state and gene expression profiles of individual cells in the brain, thereby facilitating the inference of their classification, properties, and potential physiological functions^1-6^. By targeting specific brain regions for sequencing, we can uncover the diversity of cell types within those areas^7,8^ Moreover, applying single-cell sequencing across different brain regions and species helps us understand variations in cell type composition among various brain areas^4,5,9^. Current research has identified thousands of distinct cell types in the brain^4,5,10-13^. Notably, spatial transcriptome analysis at the single-cell level further reveals the varied distribution of neuronal cells across different brain regions, significantly enhancing our understanding of the brain’s fine-scale spatial organization^11,14-18^. As techniques improve and data accumulate, we anticipate discovering more underlying design principles at the single-cell level across diverse brain regions.

Large-scale omics studies of the cerebral cortex have been conducted for years, evolving from microarray techniques to single-cell sequencing^12,19-22^. Encompassing various developmental stages, distinct cortical locations, and multiple species, these studies have yielded numerous significant discoveries^8,23-26^. Recent studies into the spatial transcriptome of the mouse cortical area have been particularly intriguing, uncovering a complex, spatially distributed cellular architecture^11,14,15,27,28^. Within the cortex, neuronal cell types demonstrate spatial gradient effects and engage in complex interactions^11,15,28^. Additionally, notable differences in neuronal cell composition across different cortical layers have been documented^16,29^. However, there has been limited research on the spatial transcriptome across the entire cerebral cortex of primates, particularly concerning the principles of spatial single-cell organization of neurons.

In this study, we investigated the spatial distribution of 26,771,303 neuronal cells across the entire cerebral cortex of macaques, examining their fine-scale spatial organization within different cortical layers. We uncovered a high degree of non-randomness in the distribution of neurons, resulting in the formation of numerous neuronal clusters. Strikingly, more than three-quarters of all neurons were located within these spatial clusters. We then analyzed the properties of various neuronal clusters, including their cell type compositions and the interactions between inhibitory and excitatory neuronal clusters. Additionally, we examined the association between the distribution of neuronal clusters and specific functional regions of the brain. Our findings suggest a common mechanism of cortical cell type utilization at the neuronal population level in primates, offering new insights into the micro-organizational principles of the primate cerebral cortex.

## Results

### Neurons in the primate cerebral cortex cluster together within each layer

Previous research has demonstrated that the ratio of excitatory (E) to inhibitory (I) neurons, known as the E-I ratio, is balanced in the layer 2/3 of the mouse cortex ^30-33^. However, the status of E-I balance in the cortex of primate brains remains elusive. In this study, we analyzed the distribution of a total of 26,771,303 neurons across six layers (layers 1-6) in the macaque brain cortex, encompassing 143 brain areas. We calculated the proportion of excitatory neurons to inhibitory neurons in each layer of every brain region. The results, as illustrated in Fig. 1a-f, indicate that, except for layer 1, the E-I ratio in layers 2 through 6 generally remains balanced across different brain areas. Moreover, with an increase in the number of excitatory neurons, there is a gradual increase in the number of inhibitory neurons, demonstrating a highly significant correlation between the two (R > 0.98, P < 1 × 10^−100^, Supplementary Data Fig. 1a). We observed that layer 1 contains more inhibitory neurons, resulting in a lower ratio of excitatory neurons to inhibitory neurons compared to other layers (Fig. 1a). Interestingly, 13 out of 33 prefrontal regions exhibit a higher number of inhibitory than excitatory neurons in layer 1. Moreover, we observed a significant increase in the E-I ratio from layer 1 to layer 6 (Supplementary Data Fig. 1b, P < 0.05), suggesting that in the deeper cortical layers of the primate brain, the proportion of excitatory neurons gradually increases.

**Fig. 1.**
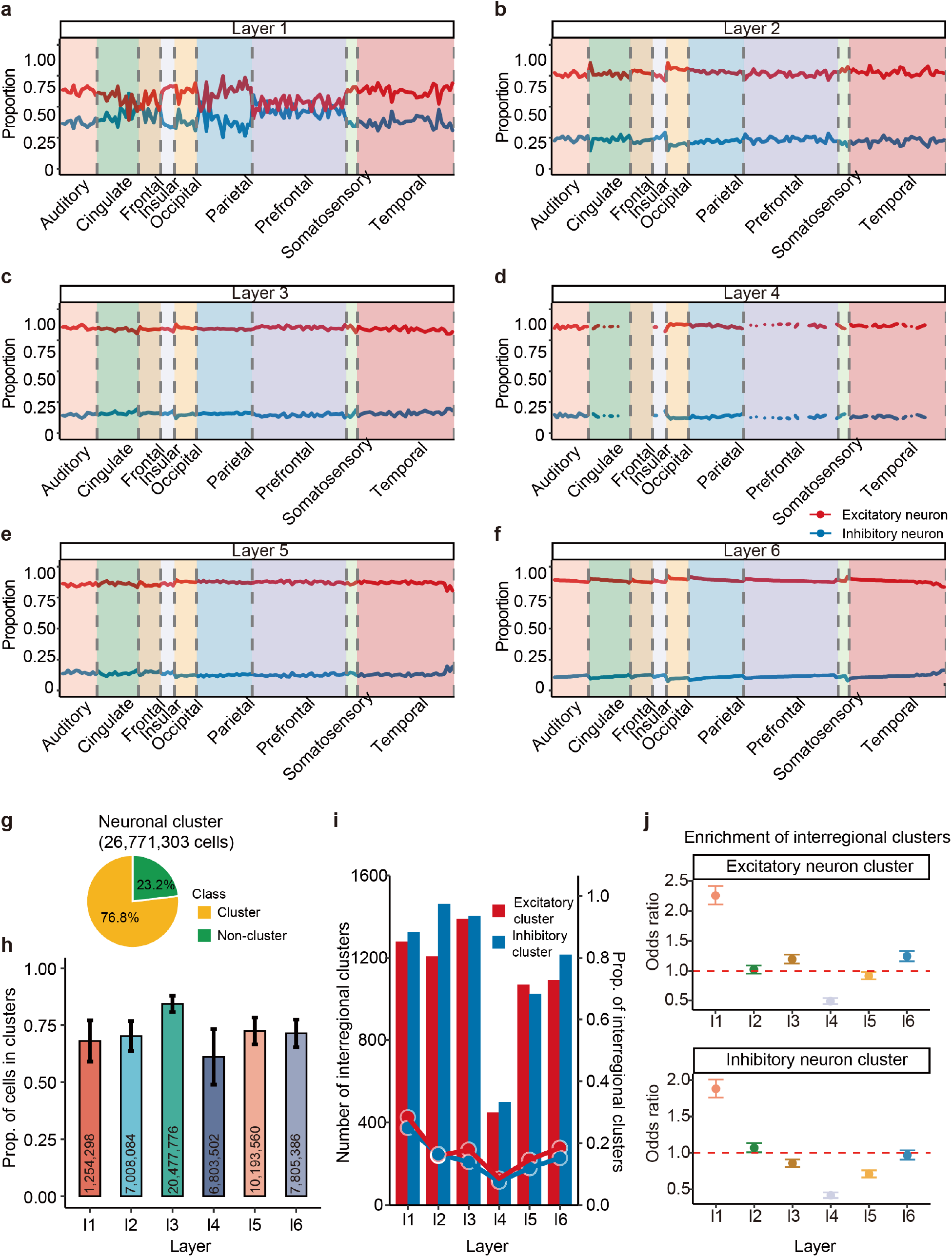
Non-random distribution of neuron cells within cortical layers of macaques. (**a-f**) Proportions of excitatory neurons (red) and inhibitory neurons (blue) in each layer (Layer 1-6) of 9 major cortical lobes (auditory, cingulate, frontal, occipital, parietal, prefrontal, somatosensory, and temporal). The y-axis quantifies their proportions, and the x-axis categorizes the cortical lobes, with dashed lines indicating regional boundaries. (**g**) The classification of all neurons shows 76.8% in neuronal clusters and 23.2% not in clusters, with a total of 26,711,303 neuron cells analyzed. (**h**) Proportion of cells in neuronal clusters for layers 1 to 6, showing that a significant number of cells in each layer are part of clusters. The error bars represent the standard error of the mean across 119 Stereo-seq chips, and the numbers indicate the total neurons analyzed in each layer. (**i**) Interregional clusters exist widely. Bar plot showing the number of interregional clusters detected in each layer, and a line plot showing the proportion of interregional neuronal clusters relative to total clusters in each layer. (**j**) Enrichment of interregional clusters in different layers. The y-axis indicates the odds ratio, with error bars representing the 95% confidence interval.

Although the ratio of excitatory to inhibitory neurons is generally balanced across different brain regions, the possibility of small-scale spatial clustering of excitatory or inhibitory neurons within layers still exists. Such localized clustering could suggest that neuronal populations in these areas perform specialized functions^34-40^. To investigate the non-randomness in the spatial distribution of neuronal cells, we employed a scanning methodology across each layer using a 1000μm x 1000μm window with a step size of 100μm, documenting the distribution of cell types within each window. We subsequently conducted 10,000 permutation experiments to evaluate the significance of the excitatory and inhibitory neuron distributions within each window. Overlapping windows that significantly enriched the same cell type (either excitatory or inhibitory) were merged into larger areas. This strategy enabled us to determine the presence of regions with significant large-scale enrichment of either excitatory or inhibitory cells and to further analyze the characteristics of these neuron clusters. Notably, even when employing a finer resolution analysis with a sliding window of 400μm x 400μm and a 100μm step size, the primary conclusions of our study remained consistent.

We observed that the size of spatial neuron clusters varied from a few cells to several thousand, with a median of 96 cells. Notably, out of the 26,771,303 neurons we examined, 20,574,158 were found within these clusters. This indicates that, overall, more than three-quarters (76.8%) of the neurons were clustered together (Fig. 1g, Supplementary Table 1). In each layer, the majority of cells were also located within clusters (Fig. 1h). The highest proportion of cells within clusters was observed in layer 3, at 84.25% (Fig. 1h), suggesting that neurons in layer 3 are more prone to clustering. Conversely, the lowest proportion of cells within clusters was found in layer 4, at 61.03% (Fig. 1h). Regionally, the occipital and frontal areas exhibited a higher tendency for clustering, particularly enriched in excitatory neurons (P = 0.012 and 0.0058, Wilcoxon rank-sum test), with rates of 78.16% and 77.37%, respectively. In clusters predominantly composed of inhibitory neurons, the cingulate area showed a higher degree of clustering, at 77.69% (P = 0.004, Wilcoxon rank-sum test).

Each of the 143 subcortical regions is spanned by at least one neuronal cluster, with 15.63% of these clusters being interregional (Fig. 1i, Supplementary Table 1). Our analysis revealed that interregional clusters typically contain more cells compared to intraregional clusters (P < 1 × 10^−100^), a trend consistent across all six layers. In clusters of excitatory neurons, inter-regional clusters are more prevalent in layer 1 (p = 1.22 × 10^−100^), layer 3 (p = 1.92 × 10^−8^), and layer 6 (p = 7.8 × 10^−10^, Fig. 1j). Conversely, in clusters of inhibitory neurons, inter-regional clusters are more likely to be found in layer 1 (p = 1.59 × 10^−73^) and layer 2 (p = 0.028, Fig. 1j). Additionally, at the lobe level, numerous clusters were found spanning two cerebral lobe areas. These findings suggest that traditional divisions of brain subregions may be reconsidered in light of fine-scale structural analyses, revealing a more complex micro-organization within the brain.

### Excitatory and inhibitory neuron clusters exhibit significant differences in properties and cell type usage

We observed significant differences between excitatory and inhibitory neuron clusters. Firstly, although inhibitory neurons constitute approximately 15% of all neuronal cells, the number of neuronal cells within inhibitory neuron clusters is similar to the number of neuronal cells within excitatory neuron clusters (10,703,572 vs. 11,964,674 cells), and this holds true for each layer (Fig. 2a, Supplementary Table 1). Thus, the presence of neuronal cells in inhibitory neuron clusters is significantly higher than expected (P < 1 × 10^−100^, Chi-square test). Additionally, the number of inhibitory neuron clusters significantly exceeds that of excitatory neuron clusters in each layer (Fig. 2a). The size of inhibitory neuron clusters is also significantly smaller than that of excitatory neuron clusters (Fig. 2b). A significant negative correlation was found between the size of excitatory neuron clusters and their E-I ratios (r = -0.28, P < 1 × 10^−100^, Fig. 2c), whereas a significant positive correlation was observed between the size of inhibitory neuron clusters and their E-I ratios (r = 0.80, P < 1 × 10^−100^, Fig. 2d). These findings suggest distinct electrophysiological properties for these two types of clusters. Moreover, the average distance between cells within excitatory neuron clusters (44.88μm +/- 22.36 μm) is significantly shorter than that within inhibitory neuron clusters (50.38μm +/- 34.50 μm, P = 1.1 × 10^−123^), indicating a higher neuronal density within excitatory neuron clusters. Intriguingly, we also identified 3,893 excitatory neuron clusters that did not contain any inhibitory neurons, with an average of 22 cells per cluster; and 179 inhibitory neuron clusters without any excitatory neurons (≥ 5 cells on average). These clusters may play specialized functions in the cortex.

**Fig. 2.**
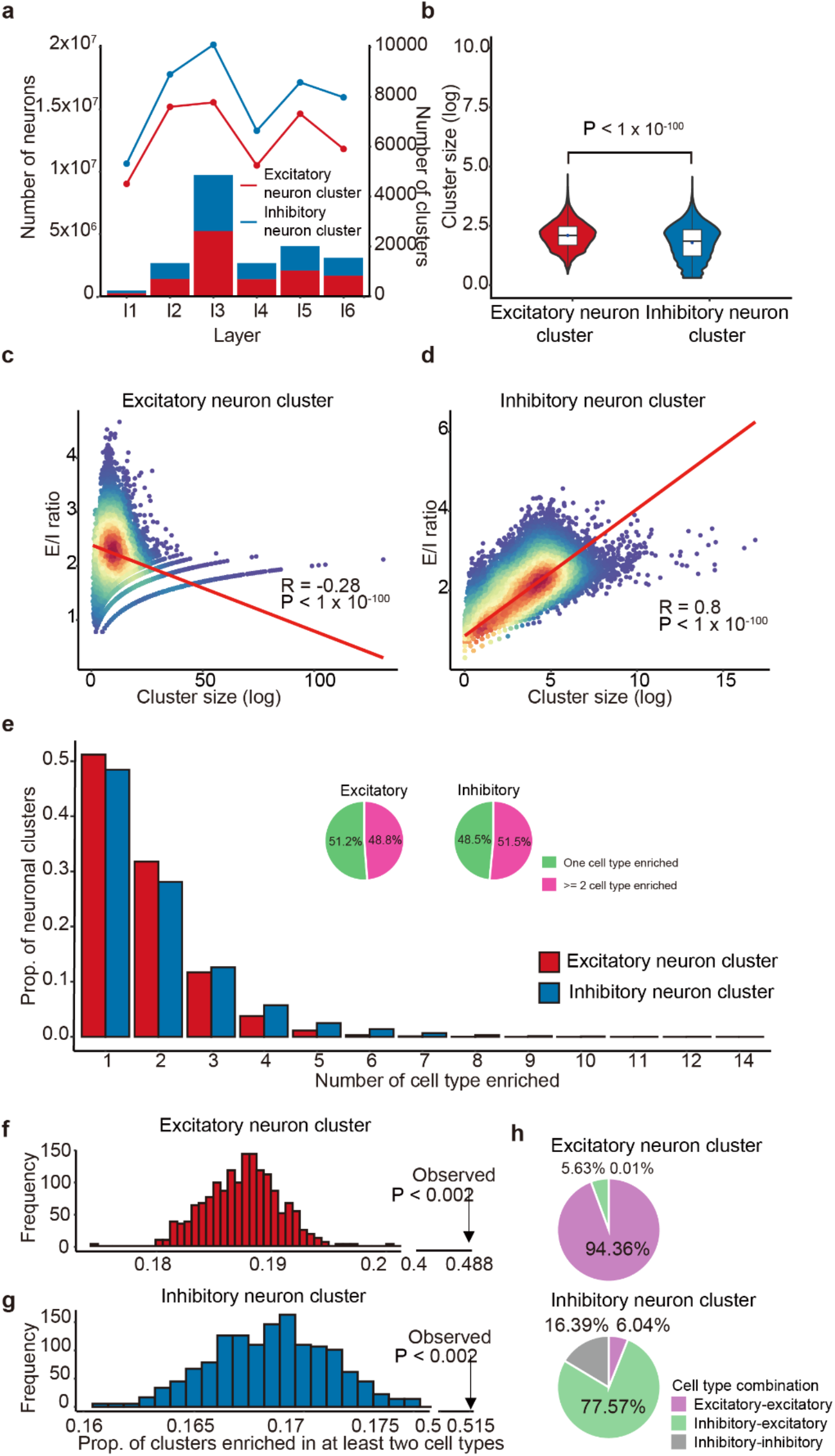
Properties and cell type usage of excitatory and inhibitory neuron clusters. (**a**) Neuron cell number and cluster distribution by cortical Layer. The bar plot illustrates that the neuron numbers in excitatory neuron clusters (red) and inhibitory neuron clusters (blue) across cortical layers 1 to 6 are similar. The line plot shows that the number of inhibitory neuron clusters (blue) is higher than that of excitatory neuron clusters (red) in each layer. (**b**) Comparison of cluster sizes of excitatory neuron clusters (red) and inhibitory neuron clusters (blue), using the Wilcoxon rank-sum test. (**c-d**) Correlation analysis of the sizes of excitatory and inhibitory neuron clusters with their E/I ratios. A scatter plot of the excitatory neuron cluster shows a negative correlation between cluster size (log scale) and E/I ratio, while the inhibitory neuron cluster exhibits a strong positive correlation between cluster size (log scale) and E/I ratio. Significance was tested using the Pearson correlation method. (**e**) The enrichment of cell types in neuronal clusters. Bar plots show the proportion of cell types enriched in excitatory and inhibitory neuron clusters. Pie charts represent the proportions of neuron clusters enriched with single or multiple cell types, indicating that nearly half of the clusters are enriched with multiple cell types. (**f-g**) Permutation test to determine the significant level of neuron clusters enriched in multiple cell types. Histograms illustrate the proportion of at least two cell types enriched in excitatory (f) or inhibitory (g) neuron clusters in the permutation experiment. These figures show that the proportion of multiple cell types enriched in a neuronal cluster is significantly higher than expected (P < 0.002). (**h**) Cell type usage in excitatory and inhibitory neuron clusters. Pie charts display the percentages of different cell type combinations within excitatory and inhibitory neuron clusters, highlighting the distinct patterns of cell-type co-occurrence.

The composition of different cell types within neuron clusters may reflect their distinct electrophysiological properties. To explore whether excitatory and inhibitory neuron clusters exhibit unique patterns of cell type usage, we first randomized the coordinates of each cell type among excitatory and inhibitory neurons and counted the number of each cell type in each cluster. This experiment was repeated 10,000 times to calculate the statistical significance of each cell type’s enrichment. Our results indicated that cell types generally tend to cluster together rather than randomly. In nearly half of the neuronal clusters (48.8% for excitatory neuron clusters and 51.5% for inhibitory neuron clusters), two or more cell subtypes significantly coexisted (Fig. 2e). These ratios are significantly higher than expected by chance, suggesting that different neuronal cell types are more likely to collaborate rather than function independently (Fig. 2f and 2g, P < 0.002). To further understand the co-occurrence among cell types, we analyzed all clusters enriched with multiple cell types. We found that more than 90% of excitatory neuron clusters consisted of combinations of excitatory-excitatory cells (Fig. 2h), while over 75% of inhibitory neuron clusters were composed of inhibitory-excitatory cell types (Fig. 2h, Supplementary Table 2). Similar trends were observed for clusters enriched with only two cell types, with excitatory-excitatory pairs accounting for over 95% in excitatory neuron clusters, and excitatory-inhibitory combinations comprising over 60% in inhibitory neuron clusters (Supplementary Table 2). These findings demonstrate that excitatory and inhibitory neuron clusters employ different strategies for cell type usage.

We also discovered that adjacent clusters tend to utilize similar cell types. By analyzing the similarity in cell type compositions between adjacent neuron clusters of the same type and comparing these with non-adjacent neuron clusters within the same layer, we observed that both excitatory and inhibitory neuron clusters exhibited significantly higher similarity among adjacent clusters than among non-adjacent ones (Supplementary Data Fig. 2a, P = 2.02 × 10^−198^ and 4.4 × 10^−143^, respectively). This observation was also corroborated by analyses using both the Pearson and Spearman correlation methods (Supplementary Data Fig. 2b-c, P < 1 × 10^−40^ in all the analysis). The pronounced similarity in cell type usage between neighboring clusters may facilitate enhanced communication among them, potentially influencing overall neural network dynamics.

### The majority of excitatory and inhibitory neuronal clusters form partnerships

Next, we explored the spatial relationship between adjacent excitatory and inhibitory neuron clusters by analyzing the overlap of cells between them, which may indicate a unique partnership. Specifically, we found that approximately 70% of excitatory neuron clusters and 62% of inhibitory neuron clusters are partnered (Fig. 3a). This ratio remains high across all layers (Fig. 3a, Supplementary Data Fig. 3). Between these partners, there’s a median of 21 shared neurons. However, the overlap of cells between them is generally low, typically encompassing less than 50% of the total cells within any specific cluster (Fig. 3b and Supplementary Table 3). Inhibitory neuron clusters tend to have more partners than excitatory neuron clusters (P = 0.018, Fig. 3c). Both types of clusters exhibit a significant positive correlation between cluster size and the number of partners (R > 0.5, p < 1 × 10^−100^, Fig. 3d and 3e), but there is no significant relationship between the sizes of the partnering clusters. These results suggest that larger neuronal populations tend to maintain balanced relationships through more partners, facilitating diverse interactions within the neural network.

**Fig. 3.**
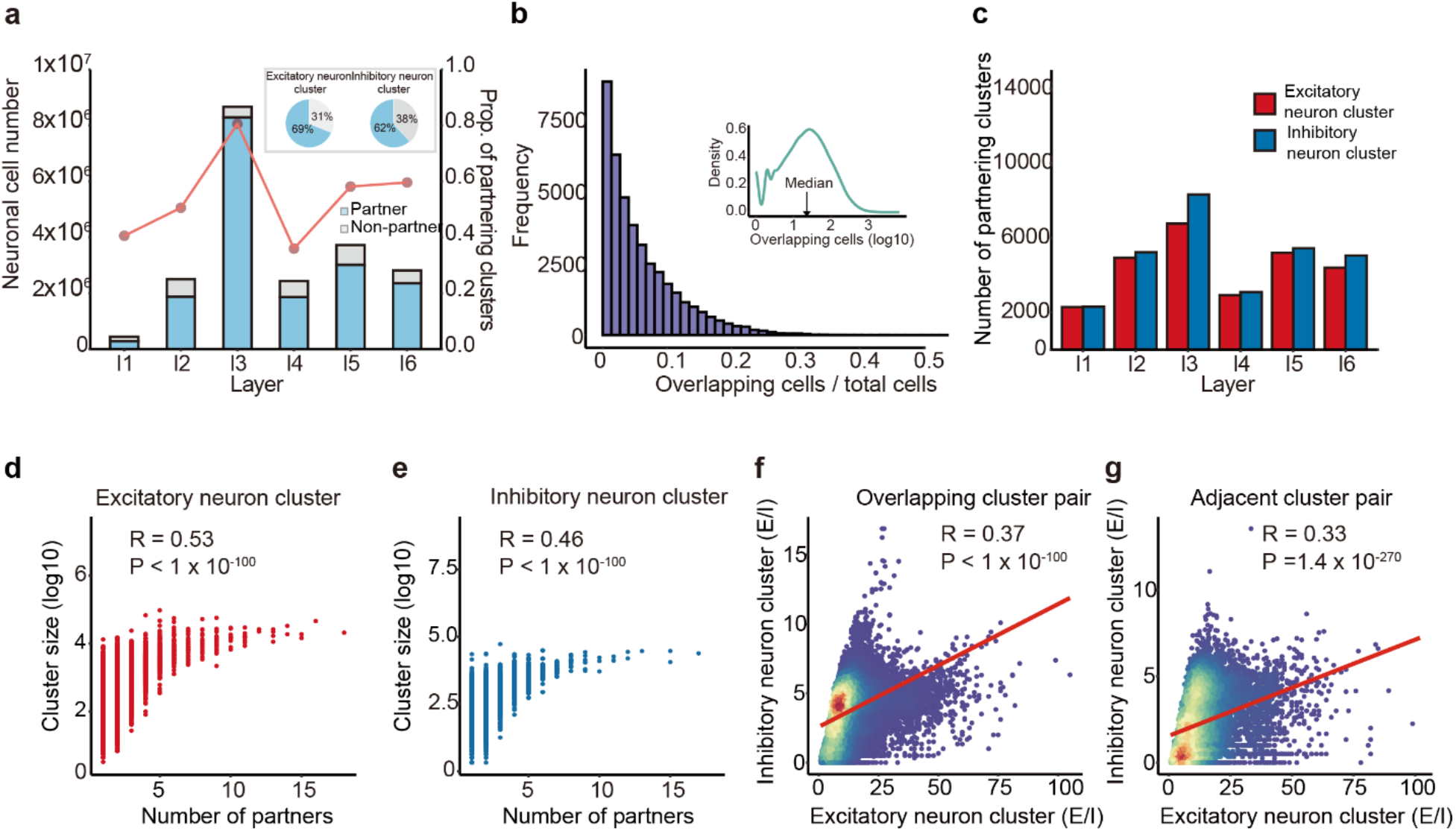
Partnership between excitatory and inhibitory neuron clusters. (**a**) The majority of excitatory and inhibitory neuron clusters have partners in each layer. A bar plot shows the number of neuronal cells in a partnership and the number of neuronal cells without a partnership. Additionally, a line plot displays the proportion of partnering clusters in each layer, and a pie chart represents the proportion of partnering clusters among all clusters for both excitatory and inhibitory neuron clusters, respectively. (**b**) The overlap between partnering clusters is relatively low. A histogram depicts the ratio of overlapping cells to total cells per partnered cluster, with a density plot showing the distribution of shared neurons. (**c**) Inhibitory neuron clusters tend to have more partners than excitatory neuron clusters. A bar plot shows the number of partner clusters for excitatory (red) and inhibitory (blue) neurons. (**d-e**) Correlation of the cluster size with the number of partners. Scatter plots for excitatory (d) and inhibitory (e) neuron clusters exhibit a strong positive correlation between cluster size (log scale) and the number of partners. Significance was tested using the Pearson correlation method. (**f-g**) The scatter plots for overlapping (f) and adjacent (g) cluster pairs reveal a similar positive correlation between the E/I ratio of the partners. Significance was tested using the Pearson correlation method.

Interestingly, we discovered a significant positive correlation between the E-I ratios of coupled excitatory and inhibitory neuron clusters (r = 0.37, P < 1 × 10^−100^, Fig. 3f). Notably, excitatory neuron clusters with low E-I ratios were more likely to form partnerships (r = -0.16, P = 1.4 × 10^−143^), whereas inhibitory neuron clusters with high E-I ratios tended to have partners (r = 0.22, P < 1 × 10^−100^). Furthermore, among the 3,893 clusters containing only excitatory neurons, significantly fewer partners were detected (p = 3.3 × 10^−156^), suggesting that these clusters tend to work independently. Even in the absence of direct overlap, the E-I ratios between adjacent excitatory-inhibitory cluster pairs showed a significant positive correlation (r = 0.33, P< 1 × 10^−100^, Fig. 3g). This suggests that the physical proximity between clusters has physiological significance. Collectively, these findings provide crucial insights into the functional organization of the neocortex at the neuronal population level.

### The distribution of neuronal clusters correlates with the physiological properties of brain regions

We explored whether the identified neuronal clusters are associated with specific physiological functions, focusing on brain regions related to the primate visual system^41^, somatosensory system^41^, and the default mode network (DMN)^42^. Initially, we observed that within the primate visual system, regions involved in early processing pathways tend to utilize larger neuronal clusters, whereas regions engaged in higher-level pathways prefer smaller clusters. This negative correlation between hierarchical level and cluster size was observed in both excitatory and inhibitory neuron clusters (Fig. 4a and 4b, R = -0.6, P < 1×10^−5^ for both datasets). Additionally, we discovered that early visual pathways exhibit a higher E-I ratio (Supplementary Data Fig. 3a-b). In the somatosensory system, excitatory neuron clusters demonstrated a similar pattern (R = -0.55, p < 0.01), further underscoring the importance of large clusters in the early stages of information processing. These findings reveal a close relationship between the distribution of neuronal clusters and the levels of information processing in which they are involved.

**Fig. 4.**
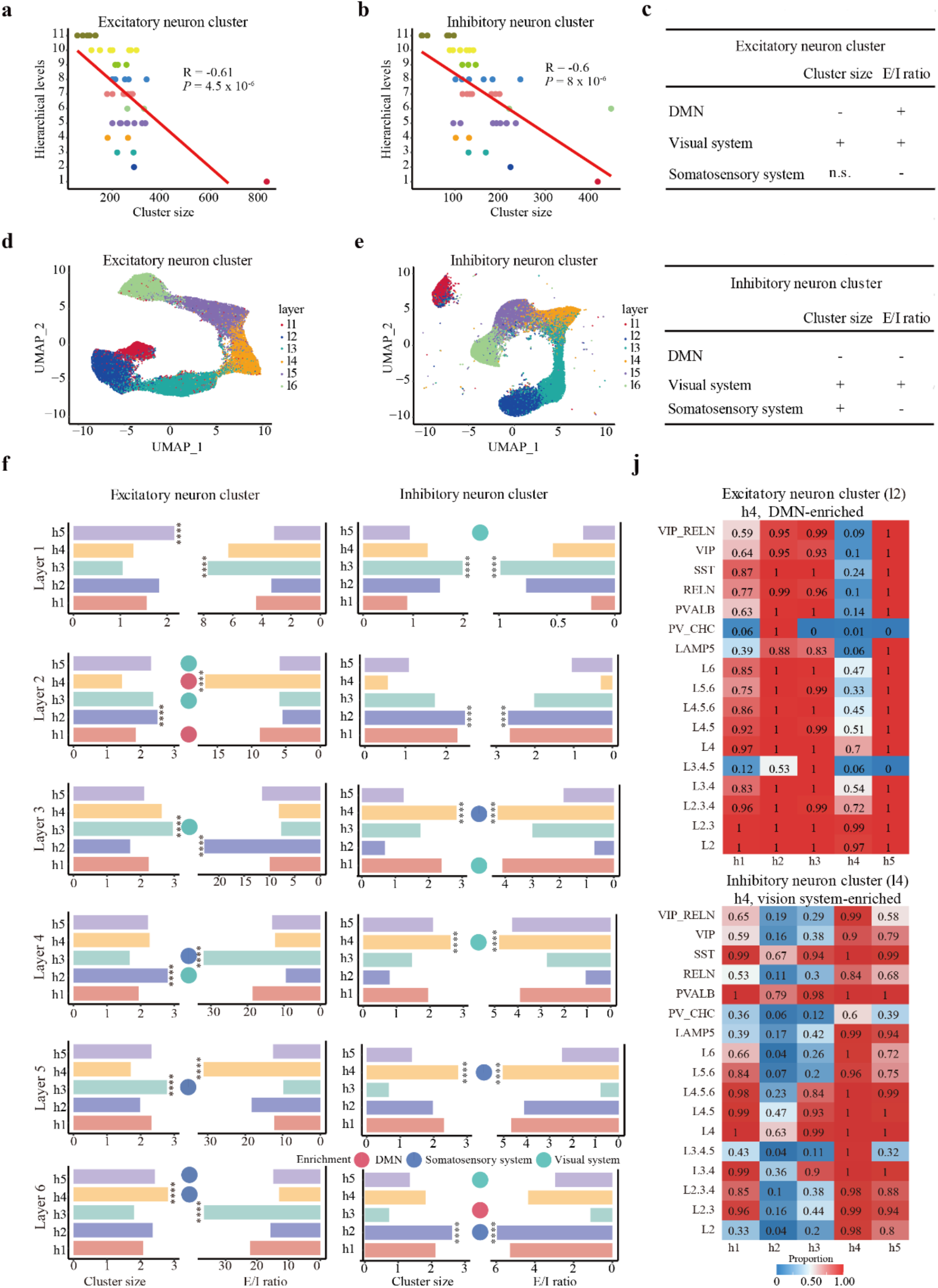
The linkage between neuronal clusters and physiological properties of brain regions. (**a-b**) Association between cluster size distribution and cortical hierarchy of the primate visual system. Scatter plots show the correlation between hierarchical levels and cluster sizes for both excitatory (**a**) and inhibitory neuron clusters (**b**), indicating that larger neuron clusters are located in early processing pathways. Significance was tested through Pearson correlation. (**c**) A table summarizes the relationship between the distribution of excitatory and inhibitory neuron clusters and different functional brain regions, including DMN, visual system, and somatosensory system, highlighting their function-specific distributions. + positive correlation with p < 0.05, -negative correlation with p < 0.05, n.s., no significant relationship (**d-e**) UMAP visualizations of excitatory neuron clusters (d) and inhibitory neuron clusters (e) display the diverse spatial distribution of these clusters across different cortical layers. (**f**) Bar plots illustrate the distribution of cluster sizes and E/I ratios across different groups, while bubble plots show their degree of enrichment in the DMN, visual, and somatosensory systems. (**j**) Distinct cell type composition for different groups. The top heatmap shows excitatory neuron clusters for layer 2, with group h4 being DMN-enriched. The bottom heatmap shows inhibitory neuron clusters for layer 4, with group h4 being visual system-enriched. In the heatmaps, values closer to 1 indicate the widespread presence of the cell type in this group.

In the brain regions associated with the DMN, both excitatory and inhibitory neuron clusters were found to be significantly smaller (p = 1.54 × 10^−17^ and 1.27 × 10^−9^, respectively, Wilcoxon rank-sum test, Fig. 4c). In contrast, within the primate visual system, the sizes of both excitatory and inhibitory neuron clusters were significantly larger than those in other brain regions (p = 3.69 × 10^−57^ and 0.00047, respectively, Wilcoxon rank-sum test, Fig. 4c). Regarding the E-I ratio, this value was generally higher in the primate visual system compared to other brain regions (p = 2.56 × 10^−26^ and 2.82 × 10^−55^, respectively, Wilcoxon rank-sum test). However, in the somatosensory system, the E-I ratio was lower than in other regions (P < 1 × 10^−5^ in both comparation, Wilcoxon rank-sum test, Fig. 4c). For the DMN, the E-I ratio of excitatory neuron clusters was higher than in other brain regions (p = 9.23 × 10^−7^), while the E-I ratio of inhibitory neuron clusters was lower than in other regions (p = 2.82 × 10^−9^, Fig. 4c). These findings indicate that different functional regions of the brain preferentially utilize neuronal clusters with distinct characteristics, reflecting the brain’s specialized strategies for processing various types of information.

Next, we investigated whether neuronal clusters with specific cell type combinations are associated with particular functional regions. Initially, we conducted a clustering analysis on the cellular composition of these neuronal clusters. Our findings revealed that both excitatory and inhibitory neuron clusters could be distinctly classified according to cortical layers 1-6 (Fig. 4d-e), demonstrating significant variations in their cell type composition across different layers. To further refine our classification of these neuronal clusters, we employed hierarchical clustering to analyze them within each individual layer.

For excitatory neuron clusters, we identified two groups associated with DMN enrichment, four groups related to the visual system and another four groups linked to the somatosensory system (adjusted P < 0.05, Fig. 4f, Supplementary Data Fig. 4). Each of these groups exhibited distinct cell type compositions. In the DMN-enriched groups, within layer 2, we identified a group, DMN-l2-Exh4, characterized by a high frequency of L2 and L2.3 cell types and minimal occurrence of L3.4.5 and inhibitory neuron cell types (Fig. 4j). This group also exhibited a relatively high E-I ratio. In layer 6, we identified a cluster, DMN-l6-Exh3, characterized by the absence of L3.4.5 cell types among excitatory neurons, with lower frequencies of L2, L2.3, L2.3.4, and L3.4 cell types, and almost no presence of inhibitory cell types. This cluster also displayed a relatively high E-I ratio. In the somatosensory system-enriched groups, within layer 5, we identified a group, SOMA-l5-Exh3, characterized by the presence of all cell types, with the highest frequency of PV_CHC cells (Supplementary Data Fig. 4a), and displaying a relatively low E-I ratio. In layer 6, we found two groups, SOMA-l6-Exh4 and SOMA-l6-Exh5, both exhibiting comparable E-I ratios and containing all types of excitatory neurons. In the vision system-enriched groups, within layer 2, we identified two clusters, VIS-l2-Exh3 and VIS-l2-Exh5, both lacking PV_CHC cells. In layer 3, we found a cluster named VIS-l3-Exh4, characterized by the absence of PV_CHC cells but the presence of other cell types (Supplementary Data Fig. 4b, Supplementary Table 4).

For inhibitory neuron clusters, we identified one group associated with DMN enrichment and four groups related to the visual system (adjusted P < 0.05, Fig. 4f, Supplementary table 4). In the visual system-enriched groups, in layer 4, we identified a group named VIS-l4-Inh4, characterized by the widespread presence of cell types and a relatively high E-I ratio (Fig. 4j). In layer 1, we found VIS-l1-Inh1, characterized by the universal presence of RELN and relatively low E-I ratios. In layer 3, we identified a group named VIS-l3-Inh1, characterized by the presence of all cell types except PV_CHC cells, with a relatively low frequency of L3.4.5 cells (Supplementary Data Fig. 4c). Within the DMN-enriched groups, at layer 6, we identified a group named DMN-l6-Inh3, characterized by a generally low frequency of most cell types and a relatively low E-I ratio (Supplementary Data Fig. 4D, Supplementary Table 4). These findings suggest that neuronal clusters with specific combinations of cell types indeed tend to appear in specific functional regions, reflecting how the brain adapts to complex information processing demands through diverse cell type configurations.

## Discussion

By analyzing the single-cell spatial transcriptome of millions of neurons across layers 1 to 6 of the cerebral cortex in macaque monkeys, we observed a highly non-random spatial distribution of these neurons within the cortical layers. Over three-quarters of the neurons were located within these clusters. Notably, different neuron cell types tend to collaborate rather than function independently. Excitatory neuron clusters and inhibitory neuron clusters follow distinct cell type combination rules; excitatory neuron clusters are primarily composed of excitatory cells, while inhibitory neuron clusters predominantly consist of a mix of excitatory and inhibitory cell types. More importantly, the majority of excitatory and inhibitory neuron clusters tend to form partnerships with each other. Finally, the spatial clustering of neurons was closely associated with the physiological functions of specific brain regions.

The spatial clustering of neurons in each cortical layer offers intriguing insights. Firstly, the total number of neurons within excitatory and inhibitory neuron clusters is roughly equivalent, indicating a balanced neuron network in this aspect^43-46^. Secondly, there is considerable variability in the cell type composition of both excitatory and inhibitory neuron clusters, with marked differences in cluster size and the E-I ratio observed. This variability suggests that each neuron cluster is context-dependent, potentially linked to unique electrophysiological functions^47,48^. Typically, excitatory neuron clusters are composed predominantly of excitatory neurons, whereas inhibitory neuron clusters feature a heterogeneous mix of both inhibitory and excitatory cell types. Consequently, much of the existing research exploring the interplay between inhibitory and excitatory neurons has predominantly focused on inhibitory clusters, examining the roles of mixed inhibitory-excitatory cell types. Further investigation into the physiological functions of purely excitatory cell types in excitatory neuron clusters is warranted. Thirdly, excitatory and inhibitory neuronal clusters may differ in their fundamental electrophysiological properties; excitatory clusters exhibit higher activity in neural signaling and effectively form fine-scale network communications^49^, whereas inhibitory clusters can shape excitatory circuits in a context-specific manner^50^. This spatial clustering may facilitate rapid synchronization across local neural networks.

The observation that the vast majority of excitatory and inhibitory neuron clusters form partnerships underscores the need for balance between excitatory and inhibitory populations at the neuronal population level^43,45,46,51^, as they collaborate to maintain the stability of neural networks^46^. This explains why larger clusters require a greater number of partners. These larger clusters may be involved in more complex information processing tasks, necessitating a more intricate balance and robust feedback through specific combinations and interactions of neuronal cells. Furthermore, the coupled excitatory and inhibitory neuron clusters, as well as adjacent clusters, are likely to co-evolve for information processing during evolution. This fine-grained structure may be crucial for accurately modeling local neuronal activity at the population level^43,44,52^.

Neurons demonstrate significant plasticity during cortical development, including experience-dependent plasticity^53-55^. We propose that this plasticity extends to spatial neuronal clusters, both developmentally and evolutionarily. Such plasticity encompasses changes in cluster size and cellular composition, which are crucial for optimizing and integrating information. Additionally, the interconnections between different neuronal clusters, including those between coupled excitatory-inhibitory clusters and adjacent clusters, may also be dynamically regulated. This plasticity illustrates how the brain responds to new information by modifying the architecture of local neural networks^56,57^, highlighting the brain’s capacity to reorganize itself in response to changing environmental demands.

Each gene possesses a unique function, despite many exhibiting high sequence similarities within the genome. Genes have also displayed signatures of variation due to genetic drift and natural selection^58-61^. Similarly, neuronal clusters may exhibit these properties. In the future, a crucial research direction will involve exploring the electrophysiological properties and functions of various spatial neuronal clusters. This raises several critical questions: Are there neuronal clusters essential for human brain cognition? Have certain neuronal clusters undergone rapid evolutionary changes? Do some neuronal clusters play significant roles in specific brain functions in primates and are responsible for particular behaviors? Ultimately, our goal is to annotate functional terms for each neuronal cluster, akin to gene functional annotations in Gene Ontology (GO) or KEGG analysis^62,63^. This is essential for advancing our understanding of the systematic mechanisms governing cell populations in the brain.

In summary, our analysis of extensive single-cell spatial transcriptome data across the entire cerebral cortex of macaques has revealed a significant non-random distribution of neural cells within cortical layers. Additionally, we observed an enrichment of diverse cell types within both excitatory and inhibitory neuron clusters, including notable partnerships between these clusters. These results indicate that the neuronal network is balanced both globally and locally. These discoveries shed light on the spatial organizational principles of neurons and their underlying mechanisms of organization across various layers, thereby providing a crucial foundation for modeling cortical neuron distributions at the single-cell level.

## Acknowledgments

We thank Dr. Robert C. Froemke and Dr. Ju Huang for helpful discussions on our results. We also thank Dr. Yidi Sun for her helpful discussions on the single-cell spatial transcriptome data of the macaque cerebral cortex and the computational support provided by the big data center of SINH. Finally, we thank all members of the Wang laboratory for their discussions. This work was supported by the National Natural Science Foundation of China (Grant Nos. 81827901 and 32170567).

## Author contributions

G-Z.W. conceived and designed the project. R.C. and G-Z.W. performed the analysis with assistance from X.N. and L.M.. R.C. and G-Z.W. wrote the paper with input from all authors.

## Competing interests

The authors declare no competing interests.

## Notes

### Competing Interest Statement

The authors have declared no competing interest.

